# GenBank 2 Sequin - a file converter preparing custom GenBank files for database submission

**DOI:** 10.1101/273441

**Authors:** Pascal Lehwark, Stephan Greiner

## Abstract

The typical wet lab user often annotates smaller sequences in the GenBank format, but resulting files are not accepted for database submission by NCBI. This makes submission of such annotations a cumbersome task. Here we present “GenBank 2 Sequin” an easy-to-use web application that converts custom annotations in the GenBank format into the NCBI direct submission format Sequin. Additionally, the program generates a “five-column, tab-delimited feature table” and a FASTA file required for submission through BankIt or for the update an existing GenBank entry. We specifically developed “GenBank 2 Sequin” for the regular wet lab researcher with strong focus on user-friendliness and flexibility. It is equipped with an intuitive graphical interface and a comprehensive documentation. The application can be employed to prepare any GenBank file for database submission. It is freely available online at https://chlorobox.mpimp-golm.mpg.de/GenBank2Sequin.html.

## Background

The typical wet lab user often annotates smaller sequences such plasmids with commercial sequence visualization and annotation software like Vector NTI Advance (Life Technologies, Invitrogen, Carlsbad, CA, USA) or Lasergene SeqBuilder (DNASTAR, Madison, WI, USA). The resulting GenBank or EMBL files, however, are not accepted for submission by NCBI. NCBI itself provides the web-based tool BankIt or the stand-alone programs Sequin and tbl2asn as annotation and/or submission tools [1], but again, these programs also do not read GenBank or EMBL files. Instead, the user must provide a so-called “five-column, tab-delimited feature table” (http://www.ncbi.nlm.nih.gov/Sequin/table.html) in order to avoid time-consuming manual feature input. However, one needs to substantially familiarize with the NCBI syntax to create such an annotation table from a GenBank entry. Moreover, tbl2asn, the powerful command line program of NCBI that creates Sequin files suitable for submission, requires both, an annotation table and some computational skills.

Unfortunately, the only public browser based file converters which generate Sequin files or annotation tables from GenBank entries (gbk2sqn and gbk2tbl; developed by Andre Villegas and Paulina Konczy, Laboratory for Foodborne Zoonoses, Guelph, ON, Canada) are no longer supported [2]. A GenBank parser (gbf2tbl.pl) provided by NCBI (ftp://ftp.ncbi.nlm.nih.gov//toolbox/ncbi_tools/converters/scripts/gbf2tbl.pl) can only partially replace the two programs. Similar to our tool described below, the script produces annotation tables and FASTA files from GenBank records. These files can subsequently be read by tbl2asn to create Sequin files for direct submission. The GenBank parser, however, is not user friendly. It is only provided as a perl script and tbl2asn must be manually executed. Last, implemented features in free standalone programs such as Artemis [3], that convert GenBank files into submission formats, require installation of these additional software.

In summary, there is a strong demand for a web-based, easy-to-use file converter, which directly converts GenBank annotations into “five-column, tab-delimited feature tables” and additionally provides Sequin files for direct submission. Therefore, we developed GenBank 2 Sequin, as part of the CHLOROBOX toolkit (https://chlorobox.mpimp-golm.mpg.de), hosted and developed at the Max Planck Institute of Molecular Plant Physiology (Potsdam/Golm, Germany). The toolbox offers software applications for the analysis of (plant derived) nucleic acid and protein sequences. Another CHLOROBOX program is GeSeq, an application for a rapid and accurate annotation of organelle genomes [4]. GenBank 2 Sequin can be used to convert GeSeq’s annotation output for database submission. Nonetheless, any custom GenBank file can be prepared for NCBI submission using GenBank 2 Sequin.

## Implementation

GenBank 2 Sequin is a web application written in Java and JavaScript, and tested with the current versions of the JavaScript enabled browsers Firefox, Chrome, IE11 and Edge. It has implemented tbl2asn v25.3 (http://www.ncbi.nlm.nih.gov/genbank/tbl2asn2) for creation of Sequin files.

## Results

GenBank 2 Sequin parses the GenBank file and converts the annotation into a tab delimited annotation table (“five-column, tab-delimited feature table”). It further extracts the nucleic acid sequence information from the GenBank file and writes it, together with the mandatory source and sequence information of an NCBI record (see below), into a FASTA file. These two files can already be used for submission through BankIt or to update an existing GenBank record. To create Sequin files for direct submission, GenBank 2 Sequin invokes tbl2asn. For this, it combines annotation table, FASTA file, and additional files that contain sequence source or author submission information (see below). As an optional feature, GenBank 2 Sequin can edit or add gene product names of coding sequences (CDS), tRNAs, and/or rRNAs in or to the annotation. This might be helpful for the revision of lager genomes. Last, GenBank 2 Sequin produces several output files for quality control (Fig. 1).

**Figure 1:**
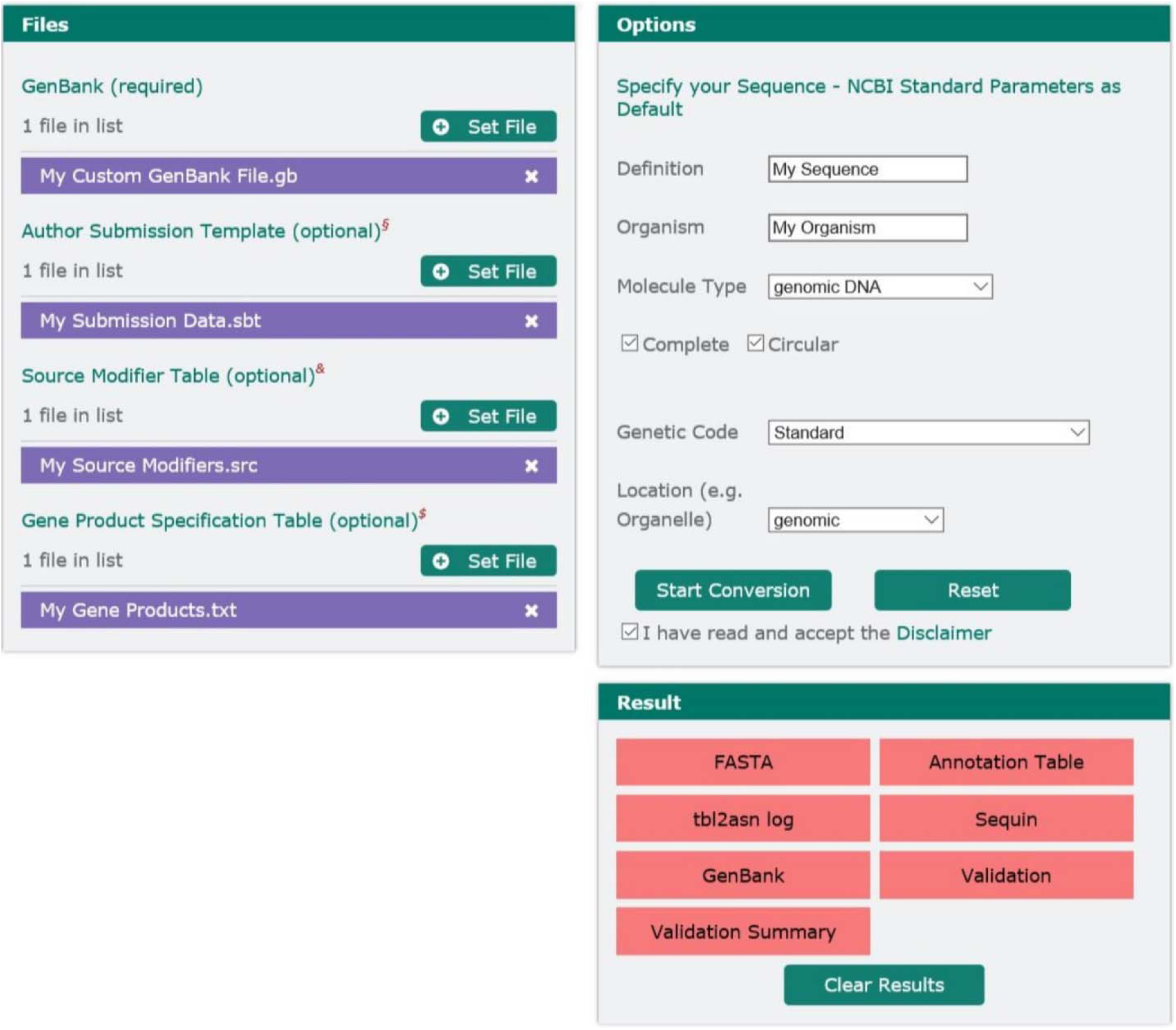
GenBank 2 Sequin Graphical User Interface. Left column (Files): File upload for the mandatory GenBank file, and optional files such as Authors Submission Template, Source Modifier Table and Gene Product Specification Table. Right column, upper box (Options): Window to specify the definition line of the GenBank entry and to add mandatory source and sequence information, such as organism or molecule type. Right column, lower box (Results): GenBank 2 Sequin output: FASTA file, annotation table, tbl2asn log, and final Sequin file for submission. For the user’s review the GenBank entry as it will be displayed later in NCBI is provided, as well as two files that include the validation of the annotation. Files can be downloaded by pressing the respective buttons. For details, see main text.

### File upload

Several file types can be uploaded to GenBank 2 Sequin:

### GenBank file

This file is mandatory and must contain the LOCUS information (either an accession number or a user defined identifier), the sequence FEATURES according to the standards of the International Nucleotide Sequence Database Collaboration (INSDC, http://www.insdc.org/documents/feature_table.html), and the ORIGIN, i.e. the nucleic acid sequence in the GenBank format. Currently GenBank 2 Sequin does not accept multi-GenBank files. Please note that FEATURES, or included qualifiers therein, which are not concise with the INSDC syntax might be modified or removed by tbl2asn (also see below). All other entries in the GenBank file, such as submitter’s information, literature references, definition line, or source information are ignored and must be provided separately by the following files or input options.

### Author Submission Template

This file contains submitter’s information and literature references, which will be later displayed in the final database entry. The template can be created at NCBI (https://submit.ncbi.nlm.nih.gov/genbank/template/submission). If no Author Submission Template is provided, GenBank 2 Sequin will use “Unknown Author” as default. Submitter’s information and literature references are also modifiable later in the Sequin file.

### Source Modifier Table

This optional upload can contain non-mandatory information for the sequence source description, such as collection site of an organism, voucher information, or a note. For controlled vocabulary see http://www.ncbi.nlm.nih.gov/Sequin/modifiers.html. The data can be either provided in the source table format *.src, (https://www.ncbi.nlm.nih.gov/genbank/tbl2asn2), or as a two-column, tab delimited text (an exemplary file can be download from the GenBank 2 Sequin web interface). Again, sequence source modifiers can also be added manually to the Sequin file prior to submission.

### Gene Product Specification Table

This table is optional as well and might be useful to revise and update lager genomes: With the help of a two column, tab-delimited text file (see example at the GenBank 2 Sequin homepage), GenBank 2 Sequin will either add or change gene product names of the annotation features CDS, tRNA or rRNA. For instance, if the Gene Product Specification Table contains the line “psaA [tab character] photosystem I P700 apoprotein A1”, GenBank 2 Sequin will search for “psaA” in the annotation. If no gene product name for “psaA” was provided in the original GenBank file (which, e.g., is the case for GeSeq output)[4], GenBank 2 Sequin will add “photosystem I P700 apoprotein A1” as gene product name. If in the original GenBank file the gene product of “psaA” was differently specified, for example as “PSI-A core protein of photosystem I”, this description will be replaced by “photosystem I P700 apoprotein A1”. If “psaA” is not present in the annotation or in the Gene Product Specification Table, no action will be taken.

### Options

In this window, the user can add/select mandatory source and sequence information, such as source organism, molecule type, location, genetic code and indicate if the sequence is complete and/or circular. In addition, the user can specify the definition line of the GenBank record.

### Output

GenBank 2 Sequin produces several output files: (i) the nucleic acid sequence in FASTA format with mandatory sequence information in the FASTA header, (ii) the annotation table, and (iii) the final Sequin file for direct submission. The first two files can be used for submission through the BankIt web interface or for the update of an existing GenBank entry.

The remaining files are for quality control: (iv) the tbl2asn log file reports any syntax errors in the original annotation as identified by tbl2asn. Those can include unknown qualifiers or feature names not allowed by NCBI. Consequently, the tbl2asn program corrects them by removing unknown qualifiers and/or changing any unknown features into misc_features (see above). (v) The annotation, as it will be displayed later in NCBI, is recorded in the GenBank output. Changes in the annotation due to potential conversion errors and/or modifications can be easily identified by comparing the user’s original GenBank file with GenBank 2 Sequin’s GenBank output using the comparison function found in many common text editors such as Microsoft Word (Microsoft Corporation, Redmond, WA, USA). (vi) Last, validation of the annotation by tbl2asn (https://www.ncbi.nlm.nih.gov/genbank/genome_validation) is provided in the files “Validation” and “Validation Summary”. Prior to submission identified annotation errors listed therein should be corrected and warnings checked. The most suitable program to correct these errors is Sequin, which also allows revalidation. Corrected files can be directly submitted.

## Conclusion

GenBank 2 Sequin is a web application that converts GenBank format annotations into the NCBI submission format Sequin. Especially for less computer-savvy wet lab researchers the tool substantially simplifies sequence submission to NCBI and hence, fills an important gap for the community. With its annotation validation by default, it also will help to improve the quality of sequence database entries.

## Authors’ contributions

GenBank 2 Sequin was conceived, designed, written and tested by PL and SG.

## Acknowledgements

We wish to thank Drs. Michel Tillich and Ralph Bock for critical reading and comments on the manuscript and an anonymous reviewer for helpful suggestions to a previous version. We are grateful to the IT Service Team of the Max Planck Institute of Molecular Plant Physiology for excellent technical assistance.

